# Characterisation of prophages in *Clostridium clostridioforme*: an understudied component of the intestinal microbiome

**DOI:** 10.1101/2024.02.29.582698

**Authors:** Suzanne Humphrey, Angeliki Marouli, Katja Thümmler, Margaret Mullin, Daniel M. Wall

## Abstract

Genome sequencing of *Clostridium clostridioforme* strain LM41 revealed the presence of an atypically high proportion of mobile genetic elements for this species, with a particularly high abundance of prophages. Bioinformatic analysis of prophage sequences sought to characterise these elements and identify prophage-linked genes contributing to enhanced fitness of the host bacteria in the dysbiotic gut. This work has identified 15 prophages, of which 4 are predicted to be intact, 2 are predicted to be defective, and 9 are unclassified. qPCR analysis revealed spontaneous release of four of the LM41 prophages into the culture supernatant, the majority of which had morphology akin to podoviruses when visualised using Transmission Electron Microscopy. We observed diversity in the lysogeny mechanisms utilised by the prophages, with examples of the classical λ-like CI/Cro system, the ICE*Bs*1 ImmR/ImmA-like system, and the Mu-like C/Ner system. Classical morons, such as toxins or immune evasion factors, were not observed. We did, however, identify a variety of genes with roles in mediating restriction modification and genetic diversity, as well as some candidate genes with potential roles in host adaptation. Despite being the most abundant entities in the intestine, there is a dearth of information about phages associated with members of the microbiome. This work begins to shed light on the contribution of these elements to the lifestyle of *C. clostridioforme* LM41.

## Introduction

The intestinal microbiome, consisting of bacteria, viruses, fungi and protozoa, plays an intimate role in contributing to the health and nutrition of its host. The microbial partners in this commensal relationship aid the host in a variety of ways, including nutrient extraction^1^, modulation of the immune and nervous systems via the activity of microbially-derived factors^2–4^, and by providing a barrier against colonisation of the gut with intestinal pathogens^5,6^. Disruption of the diversity and richness of the microbiome (known as dysbiosis) can occur through a variety of factors, including host genetics, antibiotic use, and diet and lifestyle^7–9^. Several diseases that include a dysbiosis component are associated with signature reductions and blooms of particular bacterial species^10–13^. Currently, the reasons why some species can adapt to and proliferate more readily in response to microbiome perturbation remain unclear. *Clostridium clostridioforme* (recently reclassified as *Enterocloster clostridioformis*) has been noted to proliferate rapidly in the intestines of people with a variety of conditions associated with gut dysbiosis, including obesity, type 2 diabetes and autism spectrum disorder^13–15^. Additionally, Western or high fat diets, which are established risk factors in obesity and type 2 diabetes development, significantly increase *C. clostridioforme* occurrence in the gut^16^, while the species has also been seen to increase post-antibiotic treatment^16,17^.

We recently described a novel strain of the gut commensal *C. clostridioforme*, LM41^18^. In addition to carrying multiple novel secondary metabolite biosynthetic gene clusters (BCGs), LM41 hosts several novel mobile genetic elements (MGEs), including a 192 kb plasmid, 7 integrative conjugative elements (ICEs), 5 integrative mobilisable elements (IMEs), 27 IS66 transposases, and 29 putative prophages; it is highly probable that at least some of these elements contribute to niche adaptation within the dysbiotic gut environment.

Bacteriophages (phages) are an integral facet of bacterial lifestyles and are widely distributed throughout the human gut microbiota. Estimates of the abundance of phage in the intestine are thought to approximately equal that of bacteria^19^ suggesting important roles for these viruses in regulating bacterial population dynamics, contributing to horizontal gene transfer (HGT) within and between bacterial species, and altering their bacterial hosts’ fitness in the intestinal environment. While virulent phages undergo a strictly lytic replication cycle which results in phage-mediated bacterial cell death, temperate phages can engage in a secondary lifestyle known as lysogeny. Lysogeny occurs when temperate phages integrate into the bacterial chromosome following infection of the cell, generating a prophage. The prophage remains dormant in the bacterial lysogen, replicating as part of the bacterial chromosome until an environmental signal triggers its induction into the replicative, lytic pathway. Lysogeny was historically considered to be a parasitic relationship on behalf of the phage, however, increasingly studies are demonstrating that temperate phages offer their host bacteria important advantages as trade-offs for the inherent risk associated with their carriage. Integration of a prophage into its bacterial host’s chromosome can permit expression of prophage-encoded factors that alter the host cell phenotype in a process known as lysogenic conversion. These can include virulence factors, such as the Shiga toxin carried by *E. coli* STEC Stx phages, the cholera toxin carried by the *Vibrio cholerae* CTX phage, and a variety of staphylococcal toxins carried by *S. aureus* phages and phage-inducible chromosomal islands^20,21^. Other well-characterised prophage-encoded factors include the SopE SPI-1 type 3 secretion system effector proteins of *Salmonella* Typhimurium SopEφ, and immune system evasion proteins *eib* (serum resistance; λ-like phage) in *E. coli* and *oac* (O-antigen acetylase; Sf6 phage) in *Shigella flexneri*^20^. More recently, prophages have become recognised for the protective effects that their lysogenic lifecycles can have for their host bacterial cell, with some phages encoding factors to modify their host cell’s surface in order to prevent further infection by exogenous phage, e.g. the *gp15*-encoded superinfection exclusion protein of *E. coli* phage HK97^22^, or simply by occupying attachment sites within their lysogen to prevent integration of superinfecting phage. In the latter case, expression of the CI repressor molecule by the resident phage appears to be sufficient to block the replicative cycle of infecting phages^23^, leading to destruction of the infecting phage when its ability to integrate is impeded by the resident prophage occupying the *att*C site in the bacterial chromosome. Furthermore, at the whole population level, carriage of prophages can be beneficial in enabling lysogenic communities to sample genetic material from other cells owing to the stochastic nature of phage induction^24^.

Here, we sought to gain an understanding of the biology of bacteriophages in *C. clostridioforme* LM41. We used a combination of bioinformatic and experimental approaches to reveal the genome structure, functionality, and morphology of these prophages. Our findings indicate that *C. clostridioforme* strain LM41 is poly-lysogenic for 15 prophages, the majority of which are predicted to be functional or potentially functional, and many of which carry genes with roles in facilitating restriction modification and genetic diversity, possibly contributing to the apparent proclivity of LM41 for DNA acquisition. We show that 4/15 phages are spontaneously released from LM41 under standard culture conditions, with diversity observed in their morphologies.

## Materials and Methods

### Prophage identification and annotation

For details of *C. clostridioforme* LM41 whole genome sequencing, refer to Kamat *et al* (2024). Putative prophage sequences present in the LM41 genome were identified using PHASTER^25^. Manual interrogation of PHASTER hits was performed using SnapGene Viewer software (version 5.3, www.snapgene.com) and BLASTp software (https://blast.ncbi.nlm.nih.gov), with hits deemed to be prophages or prophage remnants if they contained gene clusters conforming to one or more of the classical phage genome functional modules (lysis-lysogeny control, DNA replication, packaging and capsid, tail, and lysis). Prophage regions were annotated using Pharokka^26^ (https://usegalaxy.eu/root?tool_id=pharokka) and PhageScope^27^ (https://phagescope.deepomics.org). Pharokka parameters: Pharokka DB v.1.2.0 (downloaded at 2023-08-07 07:02:08:010437); Phanotate gene predictor; E-value threshold for mmseqs PHROGs database, 1E-05. Genome completeness assessments were performed with PhageScope. Phage regions were categorised based on completeness scores: a completeness score of 100 was categorised as ‘functional’; a completeness score of >60-<100 was categorised as ‘unknown’; and a completeness score of <60 was categorised as ‘defective’. Manual inspection of annotated genomes led us to categorise a further 5 prophages as ‘unknown’ based on unusual features predicted to affect viability (see results). For detection of putative promoter sites within lysogeny modules, PhagePromoter^28^ (Galaxy server galaxy.bio.di.uminho.pt) was used to search both strands with the following parameters: threshold, 0.5; host bacterial genus, ‘other’; phage type, ‘temperate’. Phage family (myovirus, siphovirus, podovirus) was selected for each phage as assigned in Table 2. Only hits with scores in the range 0.87-1.0 were considered. All hits are presented in Supplementary File 1.

### Bacterial strains and culture conditions

*C. clostridioforme* LM41 was grown in Fastidious Anaerobe Broth (FAB; Neogen) or on FAB agar (FAB supplemented with 1.5% agar [Formedium]) under anaerobic conditions (10% H2, 10% CO2, 80% N2, 60-70% humidity) in an A35 workstation (Don Whitely Scientific) at 37°C. All media was reduced prior to inoculation. Overnight cultures were first prepared by inoculating 5 ml of pre-reduced FAB with a single colony of LM41 from a freshly streaked plate. Fresh, pre-reduced FAB was subsequently inoculated 1:50 (v/v) with the overnight culture and allowed to grow for up to 24h under anaerobic conditions.

### Induction of *C. clostridioforme* LM41 prophages using DNA-damaging antibiotics

*C. clostridioforme* LM41 was diluted 1:50 from an overnight culture into 50 ml FAB and grown to an optical density via absorbance at 600 nm (OD600) of 0.7 under anaerobic conditions at 37°C. Cultures were induced by addition of DNA-damaging antibiotics to a final concentration of 3 μg/ml: mitomycin C (Sigma), norfloxacin (Sigma) or ciprofloxacin (Sigma). An uninduced control culture was also included. All cultures were grown for a further 16-18 h post-induction. Cultures were then centrifuged at 2800 x *g* for 30 min and the supernatants were filtered through 0.22 μm filters to remove remaining bacterial cells.

### Extraction and quantification of encapsidated DNA from induced samples

Filtered supernatants were treated with 10 μg/ml DNase I (Sigma) and 1 μg/ml RNase A (Sigma) for 30 min at room temperature, then NaCl was added to a final concentration of 1 M. After incubation for 1 h on ice, the mixture was centrifuged at 11,000 x *g* for 10 min at 4°C and the supernatant was transferred to a fresh tube. Polyethylene glycol (PEG) 8000 was added to the supernatant at a final concentration of 10% (w/v) and the mixture was incubated overnight at 4°C. Phages were precipitated from the mixture by centrifugation at 11,000 x *g* for 20 min at 4°C, with the final pellet resuspended in 1 ml phage buffer (1 mM NaCl, 0.05 M Tris pH 7.8, 1 mM MgSO_4_, 4 mM CaC_l2_). For extraction of encapsidated DNA, each sample was subject to a further DNase I treatment (20 μg/ml) for 1 h at room temperature to degrade any non-encapsidated DNA in the sample. DNase activity was stopped by addition of 20 mM ethylenediaminetetraacetic acid (EDTA) (Sigma), with incubation for 10 min at 70°C. Capsids were then opened by addition of 50 μg/ml proteinase K (Sigma) and 1% SDS to each sample, with incubation at 55°C for 1 h, mixing at 20 min intervals. The samples were subsequently transferred to fresh microcentrifuge tubes and an equal volume of phenol-chloroform-isoamyl alcohol (25:24:1; Sigma) was added to each. Samples were mixed by vortexing followed by centrifugation at 18,000 x *g* for 5 min at 4°C to allow separation of the phases. The upper phase was transferred to a fresh microcentrifuge tube and the DNA was precipitated by addition of 0.1 volumes of 3 M sodium acetate (pH 5.2) and 2.25 volumes of ice-cold 100% ethanol at -80°C for 16-18 h. Samples were centrifuged at 18,000 x *g* for 20 min at 4°C, and the pellets were washed once with ice-cold 70% ethanol before centrifuging once more. After discarding the supernatant, the pellets were air dried prior to resuspension in 50 μl nuclease-free water. Resuspended pellets were stored at 4°C for 16-18 h to allow sufficient time for solubilisation of DNA in each sample, then the DNA was quantified using a DS-11 spectrophotometer (DeNovix Inc, Wilmington, USA).

### Detection of spontaneous LM41 phage release by qPCR

The presence of phages in the bacterial supernatant was quantified using qPCR. All oligonucleotide sequences are shown in Table 1. *C. clostridioforme* LM41 was grown for 24 h in FAB under anaerobic conditions at 37°C. The culture was centrifuged at 2800 x *g* for 30 min and the supernatant filter sterilised through a 0.22 μm syringe filter to remove bacterial cells and debris. Equal volumes of supernatant were treated with 10 μg/ml DNase I (Sigma) in DNase activating buffer (50 mM Tris-HCl, pH7.5; 10 mM MgCl_2_) to degrade non-encapsidated DNA, or with an equal volume of DNase activating buffer (without DNase I) as a control. In parallel, 200 ng of LM41 genomic DNA was digested to confirm enzymatic activity of the DNase enzyme mix (see Supplementary Figure 1). Samples were incubated at 37°C for 1 h, then heated at 85°C for 15 min to inactivate the DNase enzyme and lyse any phage capsids present to release encapsidated DNA. Samples were used immediately as template for qPCR analysis.

**Table 1:**
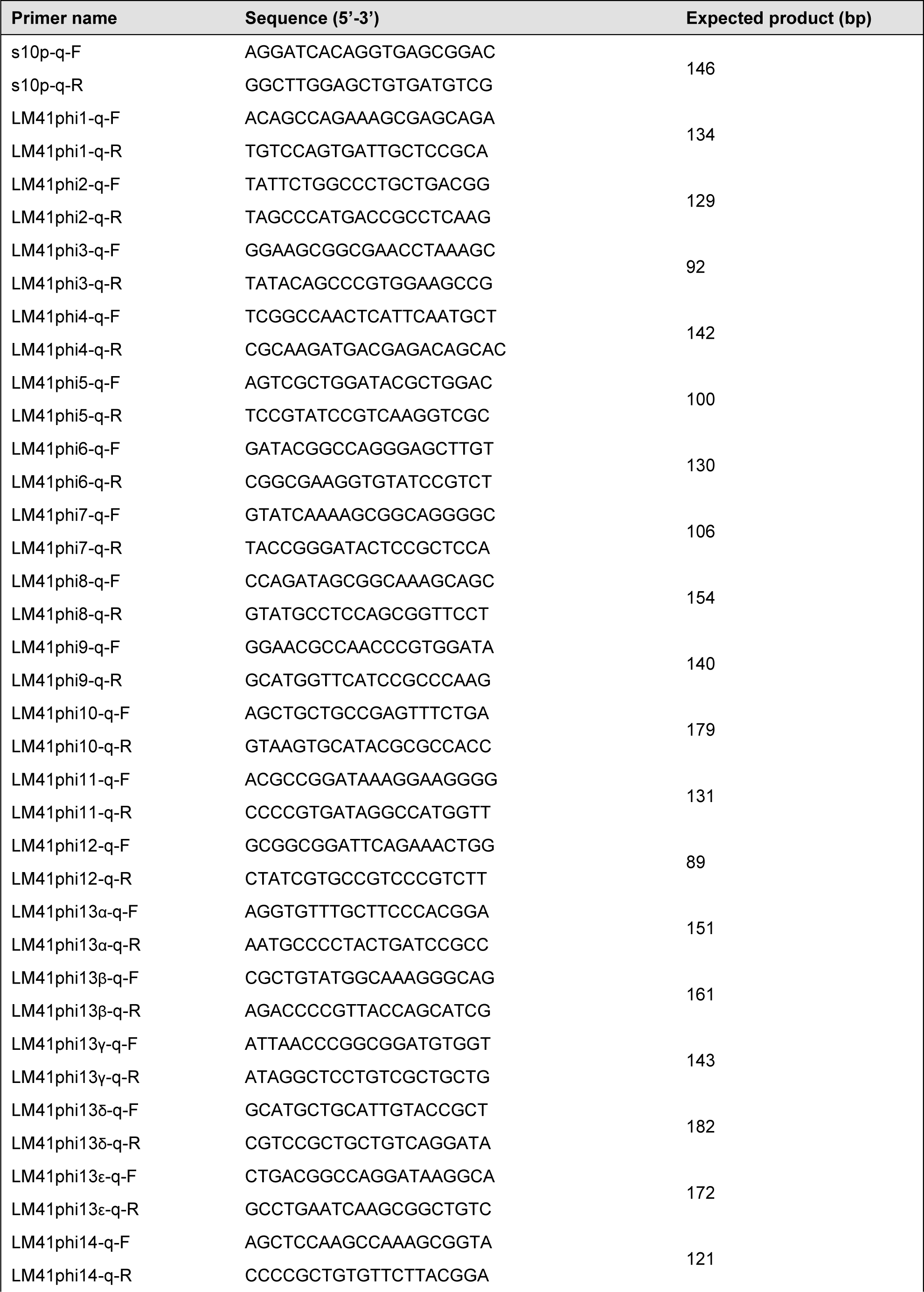
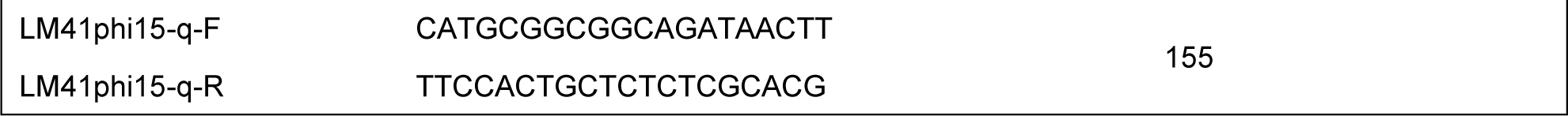
Oligonucleotides used in this study.

qPCR was performed using a CFX Connect Real-time qPCR system (Biorad). Twenty microlitre reaction mixtures were prepared using the Luna^®^ Universal qPCR Master Mix kit (New England Biolabs) as follows: 10 μl 2X Luna^®^ qPCR master mix, 7.0 μl nuclease-free H2O, 1.0 μl forward primer (final concentration 3 μM), 1.0 μl reverse primer (final concentration 3 μM), and 1.0 μl of either DNase-treated or control supernatant as template. Three technical replicates were performed per sample per primer set. Cycling conditions were: 95°C for 3 min, then 40 cycles of 95°C (10 sec), 60°C (10 sec), 65°C (30 sec). For comparison of relative quantities of phage in the sample, the 2^-ΔΔCq^ method was used, with *C. clostridioforme* LM41 small ribosomal protein 10 (*s10p*) gene used as the housekeeper.

### Purification of LM41 phages for Transmission Electron Microscopy

1.6 L of *C. clostridioforme* LM41 was grown for 24 h in FAB under anaerobic conditions at 37°C. The culture was centrifuged at 2800 x *g* for 30 min and the supernatant was filtered through a 0.22 μm filter to remove remaining bacterial cells. The supernatant was treated with 10 μg/ml DNase I for 1 h at room temperature, then NaCl was added to a final concentration of 1 M. After incubation for 1 h on ice, the mixture was centrifuged at 11,000 x *g* for 10 min at 4°C. Polyethylene glycol (PEG) 8000 was added to the supernatant at a final concentration of 10% (w/v) and the mixture was incubated overnight at 4°C. Phages were precipitated from the mixture by centrifugation at 11,000 x *g* for 20 min at 4°C, with the final pellet resuspended in 1 ml phage buffer (1 mM NaCl, 0.05 M Tris pH 7.8, 1 mM MgSO_4_, 4 mM CaCl_2_) and stored at 4°C.

### Transmission Electron Microscopy (TEM)

Carbon filmed 400 mesh copper TEM grids (AGAR Scientific) were glow discharged using a Quorum Q150TES high vacuum coater (20 mA, 30 sec). Three microlitres of precipitated phage suspensions were applied to the resulting hydrophilic carbon support films and allowed to adsorb for 3 min. Excess volume was removed by blotting and the grids were fixed for 5 min in 1 % (w/v) paraformaldehyde in phosphate-buffered saline solution. Grids were washed three times with distilled water for 30 sec, then stained with 0.5 % (w/v) uranyl formate solution for 30 sec. Sample grids were air dried at room temperature and then were examined using the JEOL 1400 FLASH TEM microscope running at 80 kV at the University of Glasgow CAF Electron Microscopy Unit (MVLS College Research Facilities). Digital images were captured at 50-150 K magnification using JEOL TEM Centre software v.1.7.26.3016 and inbuilt 2K X 2K CCD Flash camera. Particle dimension measurements were performed using ImageJ software v. 1.54h (https://imagej.net/ij/).

### Statistical analysis

Statistical analyses were performed as described in the figure legends. All analyses were performed using GraphPad Prism software version 10.1.2. Thresholds were: * *p*<0.05; ** *p*<0.01; *** *p*<0.001; *p*>0.05, not significant.

## Results

### *C. clostridioforme* LM41 is a poly-lysogen harbouring 15 prophage regions

PHASTER analysis of the LM41 genome revealed 29 predicted prophage regions^18^. These regions were subsequently interrogated to ascertain the presence of classical prophage-associated functional modules to confirm the hits as phage. Prophage completeness was estimated using Average Amino Acid Identity (AAI) comparison via PhageScope^27^, followed by manual inspection to ensure that the regions possessed essential functional modules, with genes arranged in the appropriate direction(s) to enable expression as operons. Using these criteria, 15 prophage regions were identified (Figures 1 and 2; Table 2). The genome organisation, size, and gene synteny of the majority of these prophages is reminiscent of prophages from other Gram positive species, in particular, the siphoviruses of *Staphylococcus aureus*. Four prophages (LM41φ1, φ4, φ10 and φ15) returned completeness scores of 100, suggesting that they are intact, and putatively functional. Interestingly, prophages LM41φ1 and LM41φ4 are highly similar to one another (99.02% ID across 74% of the phage sequence), with divergence occurring principally in the integrase, *cI/cro* lysogeny region, and in the latter half of the replication module.

**Figure 1:**
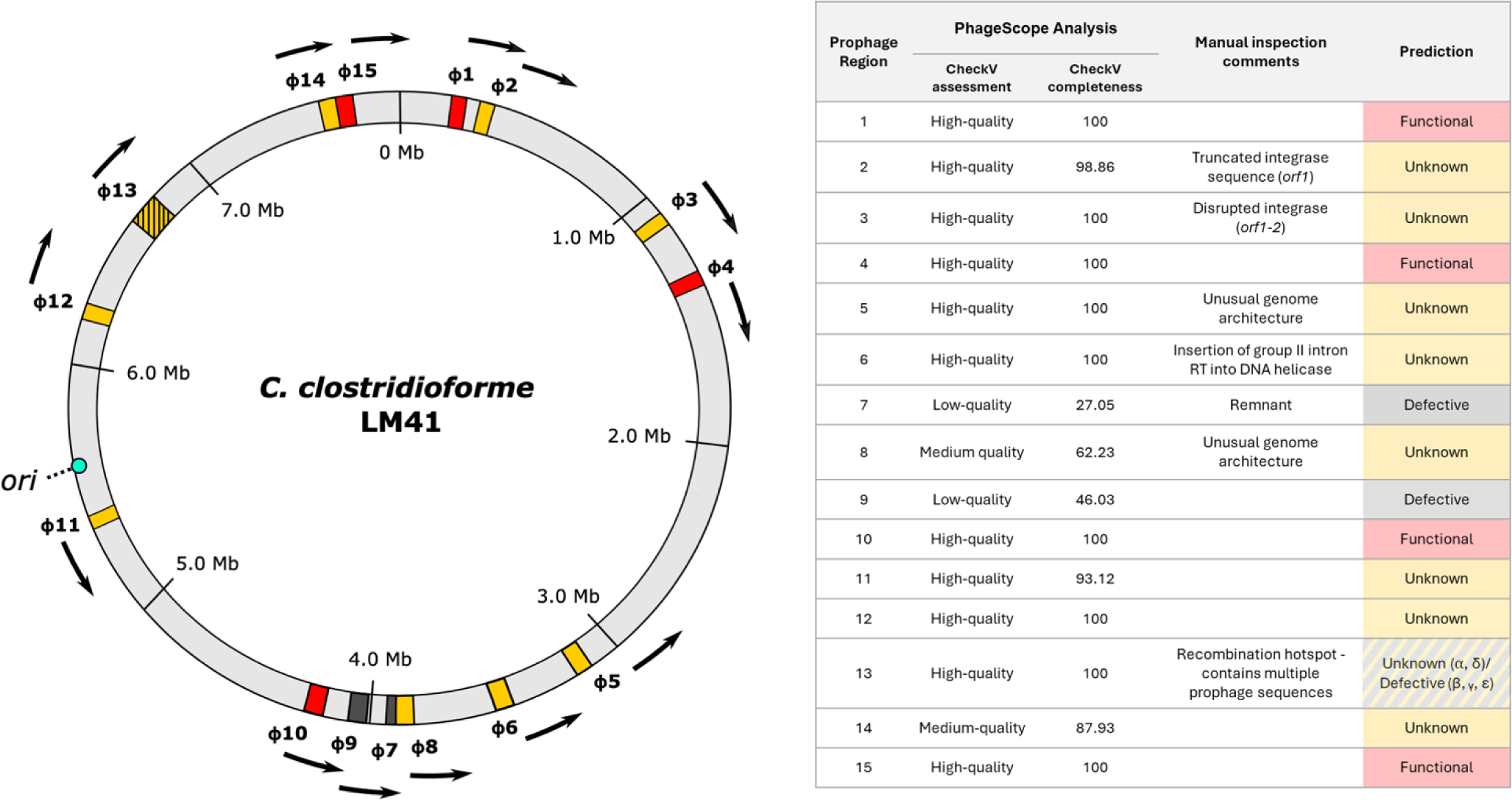
Arrangement of prophages in the *C. clostridioforme* LM41 chromosome. Locations of each of the prophage regions were mapped according to their coordinates in the LM41 chromosome. Coloured areas indicate the presence of prophages predicted to be functional (red), defective (black), or unknown (yellow) using PhageScope genome completeness assessment followed by manual inspection. Region 13 (hatched) is a putative hot-spot for phage recombination and is predicted to contain multiple phage sequences, some of which are defective remnants and some which are functionally unclassified. Arrows indicate the predicted direction of phage packaging. The bacterial chromosome origin of replication (*ori*) is indicated by the turquoise circle. Image generated using Inkscape v.1

**Figure 2:**
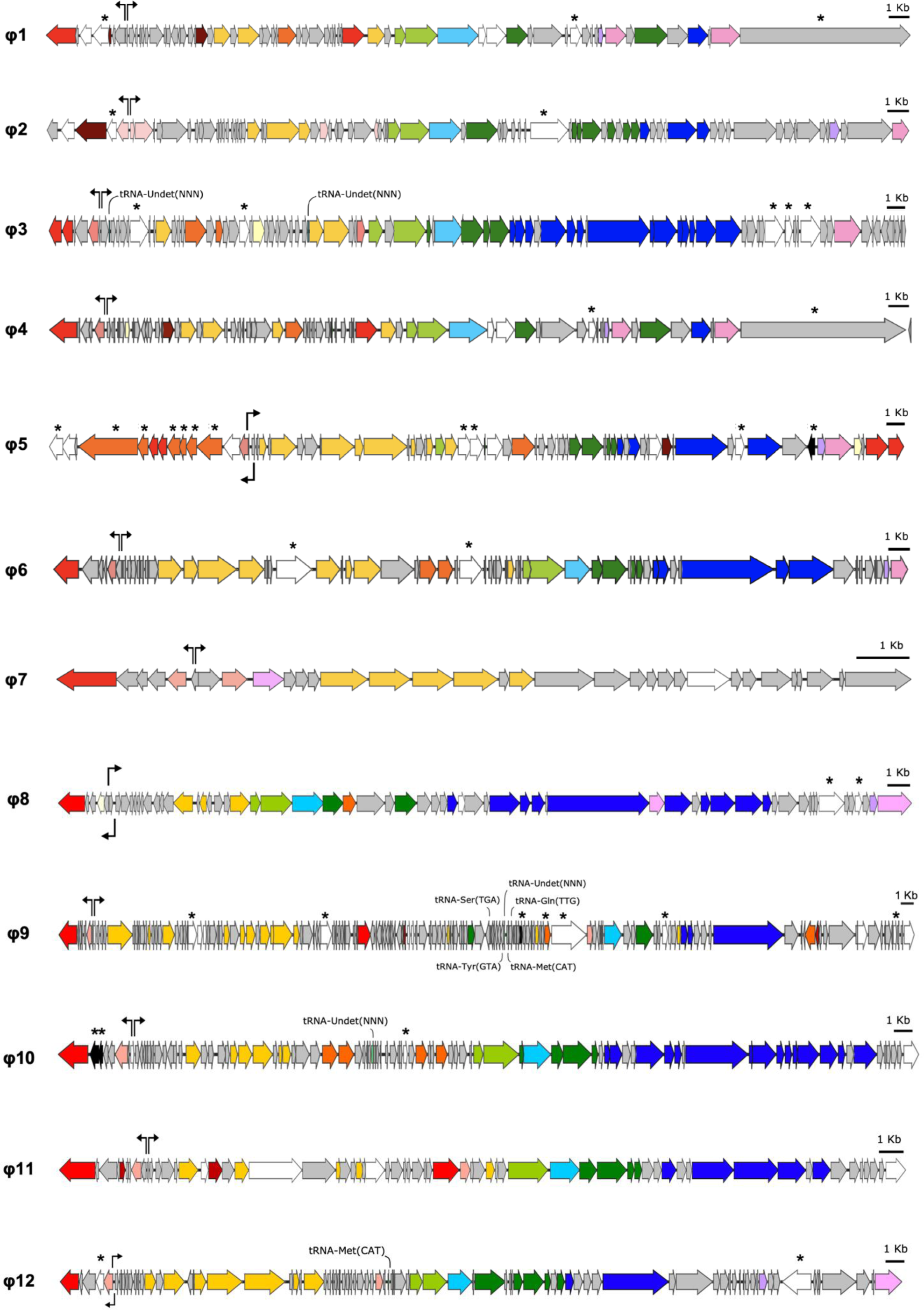
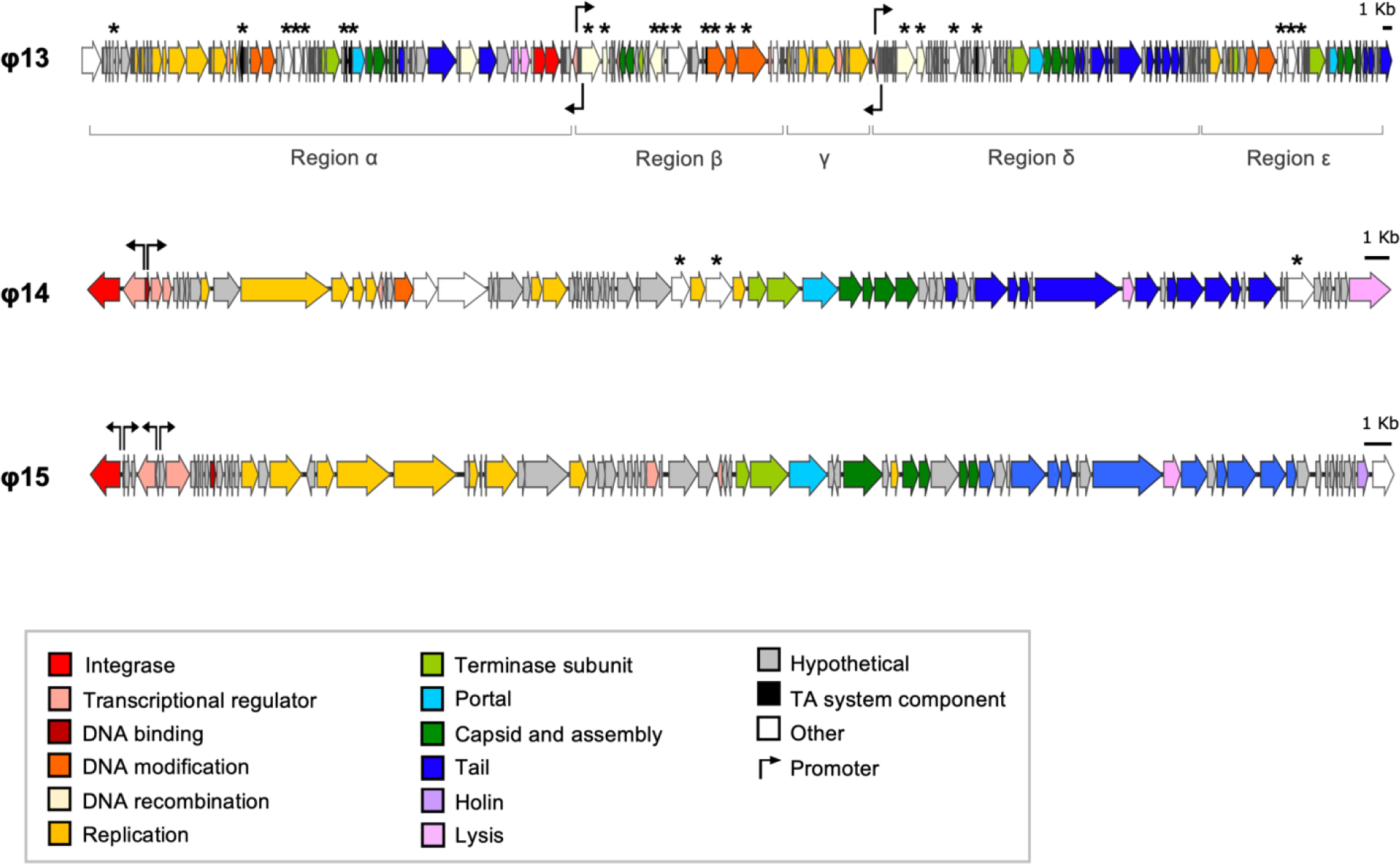
Genome maps of LM41φ1-15. Schematic maps of the ORFs predicted for each of the prophage regions identified in *C. clostridioforme* LM41. Genes are coloured according to their predicted functional group with tRNAs indicated. Putative promoters associated with lysogeny control, identified using PhagePromoter, are indicated by black arrows. Asterisks denote the presence of potential accessory genes of interest. 1 Kb scale is denoted by the black bars. Images were generated using Snapgene v.6.1.1 and InkScape v.1 software.

**Table 2:** *C. clostridioforme* LM41 prophage region characteristics.

| Phage | Location in LM41 genome | Genome size (bp) | DNA strand | Tail features |  |  |  | Closest relative (BLASTn) |  |  |  |
| --- | --- | --- | --- | --- | --- | --- | --- | --- | --- | --- | --- |
|  |  |  |  | Baseplate wedge subunit | Tail tape Measure | Sheath | Prediction | Description | Identity (%) | Cover (%) | Accession |
| φ1 | 194,517 - 235,362 | 40,846 | Forward | Orf61 | ND | ND | Podovirus | Caudoviricetes sp. isolate ctRvb1, partial genome | 94.43 | 37 | BK050138.1 |
| φ2 | 301,574 - 342,165 | 40,592 | Forward | ND | Orf65, Orf66 | ND | Siphovirus | Caudoviricetes sp. isolate ctdym5, partial genome | 95.39 | 40 | BK055266.1 |
| φ3 | 1,072,300 - 1,120,599 | 48,300 | Forward | Orf64 | Orf59 | Orf55 | Myovirus | Enterocloster bolteae strain CBBP-2 | 97.5 | 95 | CP053229.1 |
| φ4 | 1,281,572 - 1,323,457 | 41,690 | Forward | Orf63 | ND | ND | Podovirus | LM41φ1 | 99.02 | 74 | N/A |
| φ5 | 3,073,910 - 3,122,116 | 48,207 | Reverse | ND | Orf58 | ND | Siphovirus | Blautia pseudococcoides strain SCSK | 79.12 | 30 | CP053228.1 |
| φ6 | 3,448,526 - 3,490,199 | 41,674 | Reverse | ND | Orf52 | ND | Siphovirus | Caudoviricetes sp. isolate cthCz6, partial genome | 93.24 | 38 | BK022268.1 |
| φ7 | 3,885,600 - 3,901,899 | 16,300 | Reverse | ND | ND | ND | - | Caudoviricetes sp. isolate ct1dl13, partial genome | 78.11 | 58 | BK021065.1 |
| φ8 | 3,846,360 - 3,885,591 | 39,232 | Reverse | Orf47, Or48 | ND | Orf39 | Myovirus | Caudoviricetes sp. isolate ctELP1 | 79.38 | 65 | BK049563.1 |
| φ9 | 3,997,064 - 4,063,629 | 66,566 | Reverse | ND | Orf133 | ND | Siphovirus | Caudoviricetes sp. isolate ctRyL7, partial genome | 91.04 | 68 | BK049246.1 |
| φ10 | 4,120,662 - 4,165,801 | 45,140 | Reverse | Orf80, Orf81 | Orf75 | Orf71 | Myovirus | Caudovirales sp. isolate ctjl31, partial genome | 93.81 | 71 | BK029493.1 |
| φ11 | 5,193,058 - 5,226,919 | 33,862 | Reverse | ND | Orf53 | ND | Siphovirus | Caudoviricetes sp. isolate ctpXm10, partial genome | 92.98 | 48 | BK024821.1 |
| φ12 | 6,190,004 - 6,237,536 | 47,533 | Forward | ND | Orf63 | ND | Siphovirus | Enterocloster bolteae strain ATCC BAA-613 chromosome | 89.09 | 39 | CP022464.2 |
| φ13 | 6,632,999 - 6,769,447 | 136,449 | Forward |  |  |  |  | N/A | N/A | N/A | N/A |
| $\alpha$ | 6,632,999 - 6,684,039 | 51,041 | Forward | ND | Orf65 | ND | Siphovirus | Siphoviridae sp. isolate ctgM31, partial genome | 92.9 | 75 | BK028648.1 |
| $\beta$ | 6,684,149 - 6,706,426 | 22,278 | Forward | ND | ND | ND | - | Caudoviricetes sp. isolate ct1iX6, partial genome | 81.66 | 26 | BK023115.1 |
| $\gamma$ | 6,706,413 - 6,714,990 | 8,578 | Forward | ND | ND | ND | - | Lachnoclostridium sp. YL32 chromosome, complete genome | 95.82 | 96 | CP015399.2 |
| $\delta$ | 6,715,266 - 6,749,468 | 34,203 | Forward | Orf174 | Orf184, Orf185 | Orf180 | Myovirus | Lachnoclostridium sp. YL32 chromosome, complete genome | 97.62 | 53 | CP015399.2 |
| $\epsilon$ | 6,750,116 - 6,769,447 | 19,332 | Forward | ND | Orf227 | ND | Siphovirus | Siphoviridae sp. ctquf9, partial genome | 86.92 | 88 | BK014815.1 |
| $\phi 14$ | 7,473,893 - 7,529,677 | 55,785 | Forward | Orf65 | Orf61 | Orf57 | Myovirus | No matches with coverage > 1% | N/A | N/A | N/A |
| $\phi 15$ | 7,532,147 - 7,579,680 | 47,534 | Forward | Orf72 | Orf68 | Orf63 | Myovirus | Lachnoclostridium sp. YL32 chromosome, complete genome | 90.62 | 46 | CP015399.2 |

ND, Not detected

Prophage regions LM41φ7 and LM41φ9 returned poor completeness scores (<60), suggesting that these are defective phages due to loss of parts of their coding sequence. Indeed, LM41φ7 appears to be a prophage remnant, encoding only lysogeny and replication modules before being interrupted by the integrase of LM41φ8.

Regions LM41φ2, φ8, φ11 and φ14 were classified as ‘unknown’ on the basis that they returned completeness scores of >60-<100. In addition, though prophages LM41φ3, φ5, φ6, φ12 and φ13 returned completeness scores of 100, manual inspection of their genomes revealed unusual characteristics predicted to affect prophage viability, hence we also categorised them as ‘unknown’ pending further investigation.

LM41φ3 exhibits peculiarities in its integrase (*orfs1-2*); namely, that *orf2* carries a putative frameshift mutation that results in a premature stop codon after residue P199, splitting the integrase sequence, and likely rendering the prophage defective. Meanwhile, compared to the other prophages, LM41φ5, has an unusual genomic composition, with a larger lysogeny module (*orf1-13*) containing two putative integrase sequences (*orf6-7*) located between proteins comprising a complete type I restriction modification system. In the case of Orf6-7, these putative integrase proteins are smaller than expected for a typical functional integrase (Orf6, 165 aa; Orf7, 167 aa) suggesting that these may be an integrase truncated by mutation.

We attempted to investigate this by searching for related sequences to determine whether a premature stop codon had been introduced by single nucleotide polymorphism to φ5, however a BLASTn search of the sequence encompassing *orf6-7* returned no matches with significant similarity, leaving us unable to resolve this question at this time. φ5 also carries two open reading frames (ORF) predicted to encode recombinases (*orf68-69*) at the terminal end of the prophage sequence. These recombinases are closer in length to that expected for a functional integrase protein (Orf68, 420 aa; Orf69, 282 aa), raising questions as to whether one of these could catalyse the integration of the phage at the *att*B site if indeed the first integrase is non-functional. In addition, we were unable to identify certain components essential for virion packaging in LM41φ5, namely the large terminase subunit and portal protein, raising questions about the capability of this phage to package its genome into procapsids.

LM41φ6 and LM41φ12 exhibit anomalies in their DNA replication and lysis modules, respectively, on account of insertion of a putative group II intron reverse transcriptase/maturase (LM41φ6 *orf20*; LM41φ12, *orf78*). LM41φ6 *orf20* is positioned between two putative DNA helicases (*orf17* and *orf21*). *orf17* and *orf21* are not duplicated genes as they do not display any significant nucleotide similarity. Rather, BLASTn analysis of *orf17* and *orf21* sequences revealed hits with high similarity to the virulence associated protein E (putative DNA helicase) protein from Caudoviricetes sp. isolate ctdym5 [Acc: BK055266] (*orf17*: 99.92% identity, 51% cover, E value 0.0; *orf21*: 99.83% identity, 47% cover, E value 0.0), suggesting that *orfs17* and *21* are one ORF that has been split by the insertion of the putative group II intron. This likely eliminates the functionality of the helicase protein and renders the phage incapable of initiating replication following activation. In LM41φ12, *orf78* is positioned divergently to its flanking genes, potentially affecting transcription of the late module as a polycistronic transcript and therefore affecting the ability of the phage to induce host lysis.

Finally, the data obtained for region φ13 were puzzling. A completeness score of 100 was returned for this 136.5 kb region, suggesting potential functionality, however manual inspection revealed the presence of a variety of prophage sequences with differing levels of completeness. Some of the sequences within region 13 were reminiscent of Mu-type phage, hence, we divided the 136.5 kb region into sub-regions (α-ε) by matching the predicted functions of the ORFs relative to the functional modules expected for a complete Mu-type phage (i.e. a transposase, ATP-binding or DNA replication protein indicated the likely start of a phage sequence, while a tail or recombinase indicated the likely end). We then refined the regions by searching for hits using BLASTn and assessing completeness using PhageScope (Figure 2).

Regions φ13β, γ and ε returned low completeness scores (49.4, 18.8 and 41.2, respectively), suggesting that these areas are defective phage remnants. The most complete stretches of prophage genome in this region are the 51.0 kb φ13α (*orf1-79,* completeness score 99.65) and 34.2 kb φ13δ (*orf128-192,* completeness score 94.11) regions (Figure 2). Prophage φ13α appears to encode most of the required modules for replication and assembly of phage virions. However, despite encoding two putative integrase genes at its 3’ terminus, it lacks a clear lysogeny module and does not encode a Mu transposase C-terminal domain-containing protein to permit transposable replication, suggesting that it may be incomplete. Prophage φ13δ has high similarity to *Clostridium* phage Villandry (BLASTn: 95.82% identity, 96% cover, E value 0.0, Acc ON453902.1) and is reminiscent of the prototypical transposable phage Mu, encoding putative candidates for a repressor (*orf129*), a *ner*-like transcriptional regulator (*orf139*), a Mu transposase (*orf140*), a Mor transcription activator family protein (*orf158*) and structural components such as Mu-like prophage I protein (*orf167*), Phage Mu protein F like protein (*orf166*) and Mu-like prophage major head subunit gpT (*orf169*).

### LM41 prophages exhibit diversity in their lysogeny control mechanisms

Our analysis revealed diversity in the lysogeny control mechanisms of the LM41 prophages, with three groups identified: classical λ-like CI/Cro systems^23^; ImmR/ImmA-like systems similar to that used by the *Bacillus subtilis* integrative conjugative element ICE*Bs*1^29^; and systems reminiscent of the CI/Cro-like C/Ner system of *E. coli* phage Mu^30^.

Most prophages (10/15) appear to possess lysogeny systems reminiscent of the classical λ-like CI/Cro system. Prophages LM41φ1, φ3, φ4, φ6, φ7, φ9, φ10, φ11, φ14 and φ15 each possess a pair of adjacent divergently transcribed genes in their putative lysogeny modules, several of which are predicted to be helix-turn-helix transcriptional regulators, representing probable CI-like repressors. Using PhagePromoter^28^, we detected divergent promoters in the intergenic region between these ORFs (Figure 2), lending support to our hypothesis that these ORFs encode a CI/Cro-like lysogeny switch in these phages.

Prophages LM41φ2, φ5 and φ12 appear to encode systems analogous to the ImmR/ImmA regulatory systems that have been described for *B. subtilis* ICE*Bs*1^29^ and some phage-inducible chromosomal islands in staphylococci^31^. In the lysogeny modules of φ2 and φ12, *orf*4 is predicted to encode an ImmA/IrrE family metallo-endopeptidase which is located directly adjacent to *orf5*. Orf5 is predicted to be a helix-turn-helix (HTH) transcriptional regulator, which we propose to be ImmR-like based on its functional prediction and synteny with *immR* from ICE*Bs*1. Importantly, *orfs4-5* are transcribed in the same direction (leftward), while the downstream ORF (*orf6*), predicted to encode a second HTH transcriptional regulator, is transcribed in the rightward direction towards the DNA replication module. Within the intergenic region between *orf5* and *orf6* in φ2, PhagePromoter predicted the presence of two divergent promoters akin to that observed for P*immR* and P*xis* in ICE*Bs*1 (Figure 2). Conversely, in φ5 and φ12, a pair of convergent promoters were predicted in the intergenic regions between *orfs13-14* (φ5) and *orfs5-6* (φ12), suggesting that transcriptional interference similar to that observed in coliphage 186^32^ may play a role in regulating the lysogenic-lytic control switches in these prophages.

The final lysogeny system that we observed is reminiscent of the C/Ner system of *E. coli* phage Mu. In phage Mu, the 197-aa repressor protein (C) is divergently transcribed from the 76-aa DNA binding protein, Ner, which functions to negatively regulate transcription of the replicative transposition genes^30^. LM41φ13 appears to be a highly plastic region on the LM41 chromosome that contains the multiple Mu-type prophage/remnants arranged consecutively, which we have designated φ13α-ε. In LM41φ13, Orf80, located in region φ13β, is predicted to be a 168-aa HTH transcriptional regulator and is transcribed divergently from the downstream gene, *orf*82, which is also predicted to encode a 51-aa HTH transcriptional regulator (BLASTp: 100% identity, 98% cover, E value 1e-26, Acc: WP_303009215.1) (Figure 2). Immediately downstream, *orf*83 is predicted to encode a protein with a Mu transposase C-terminal domain. The similar size, synteny and functional predictions of *orf60*, *orf62 and orf63* from LM41φ13β with that of the genes encoding C, Ner and the transposase from the classical phage Mu, suggests that the functional ancestor of this phage utilised lysogeny regulation mechanism similar to that of phage Mu. A similar arrangement is present in region φ13δ, however 9 ORFs with hypothetical functions are located between the putative *c* (*orf129*) and *ner* (*orf139*) homologs.

### LM41 prophages are predicted to be morphologically diverse

Excepting the remnants of φ7, φ13β and φ13γ, genome analysis indicated that all LM41 prophages are tailed, with each of the major tail groups represented (Table 2). Prophages φ1 and φ4 are predicted to carry a single tail gene (*orf61* and *orf63*, respectively). Their products are predicted to be baseplate wedge subunits with homology to tail fibre (spike) proteins, suggesting that these may be podoviruses. Tail sheath proteins were observed in the genomes of φ3, φ8, φ10, φ13δ, φ14 and φ15, suggesting that these may be myoviruses with contractile tails. The remaining prophages, φ2, φ5, φ6, φ9, φ11, φ12, φ13α and φ13ε, possessed tape measure proteins but no sheaths, suggesting that they may be siphoviruses.

### Phage particles are spontaneously released from *C. clostridioforme* LM41

We sought to determine whether we could identify any of the phages in the supernatant of LM41 cultures. Treatment with classical SOS-inducers such as mitomycin C, ciprofloxacin and norfloxacin failed to induce lysis of LM41 cultures and did not lead to significantly higher levels of encapsidated phage DNA in induced cultures compared with basal release in untreated cultures (Figure 3), suggesting that these chemicals are not potent inducers of LM41 prophages under the conditions tested.

**Figure 3:**
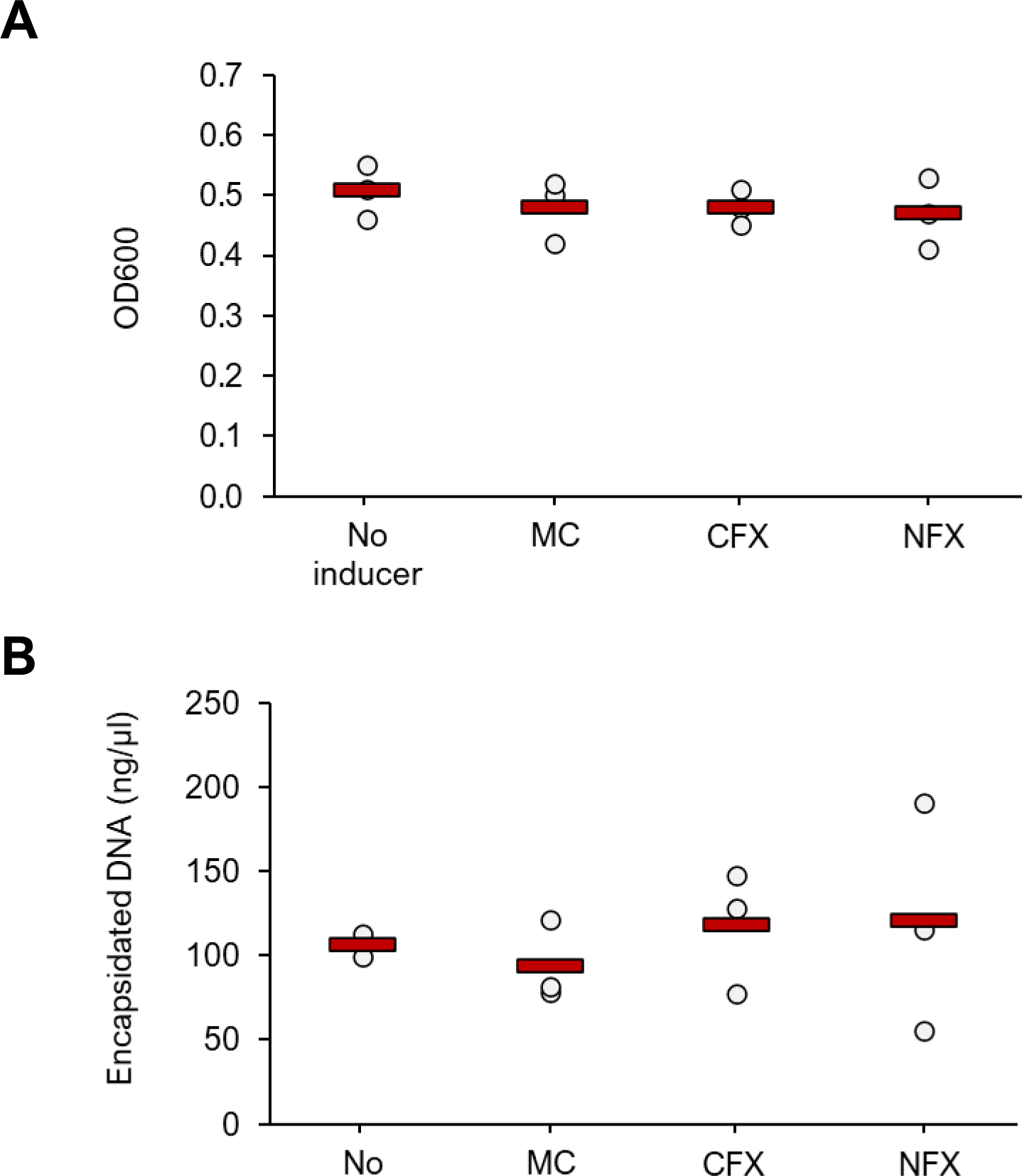
Common SOS-response inducing agents are not potent inducers of *C. clostridioforme* LM41 prophages. *C. clostridioforme* LM41 cultures were induced with common SOS-response-inducing chemicals, mitomycin C (MC), ciprofloxacin (CFX) or norfloxacin (NFX) and grown for 16-18 h at 37°C in anaerobic conditions. An uninduced control sample was also included. OD600 values were obtained for each culture after 16-18 h to determine if phage-induced lysis had occurred (**A**) and encapsidated DNA was purified and quantified for each culture (**B**). Data points are from three independent experiments (n = 3), with mean values shown as red bars. All data were tested for significance using a one-way ANOVA with Tukey post-hoc tests; no statistically significant differences were observed between the groups (*p*>0.05).

Using a dual approach, we examined the profile of phages released following spontaneous induction for our subsequent experiments. Firstly, qPCR was used to identify the presence of encapsidated DNA (indicative of phage) in LM41 culture supernatants. Briefly, filtered supernatants were divided into duplicate samples, of which one was DNase treated and the other was kept as an untreated control. DNase treatment enabled differentiation of encapsidated phage DNA, which is protected from degradation by the phage protein capsid, from DNA present in the sample that has been released from lysed bacterial cells. For all targets, DNase treatment reduced the quantity of DNA present in the sample (Figure 4A). Following treatment, levels of the bacterial small ribosomal subunit protein 10 (*s10p*) housekeeper gene were reduced below the respective no template control (NTC), suggesting comprehensive degradation of bacterial DNA in the sample. In contrast, excepting φ13ε, each of the phage targets remained detectable relative to their respective NTCs, suggesting the presence of phage particles in the supernatant, albeit often at extremely low levels. The differences in mean Cq values between treated and untreated samples were lowest for φ1 (5.11), φ4 (8.06), φ2 (8.37) and φ10 (8.69), suggesting that these phages were most abundant. Indeed, quantification of the levels of each phage in the sample relative to the *s10p* housekeeper was performed using the 2^-ΔΔCt^ method and showed that φ1 was the most abundant phage in the sample (mean ± SD: 28.45 ± 23.35), followed by φ4 (2.95 ± 0.72), φ2 (2.90 ± 1.99) and φ10 (1.87 ± 0.26) (Figure 4B).

**Figure 4:**
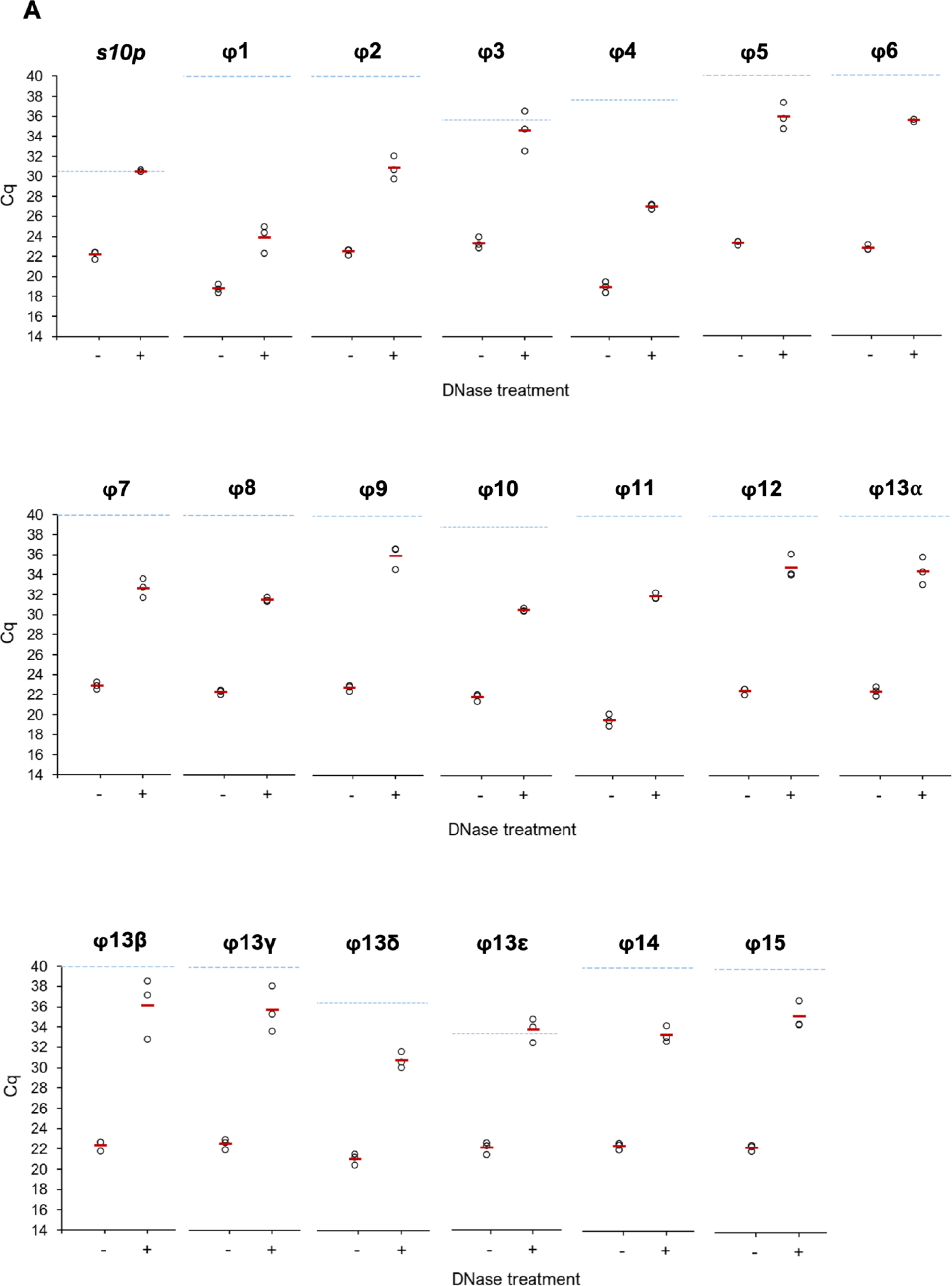
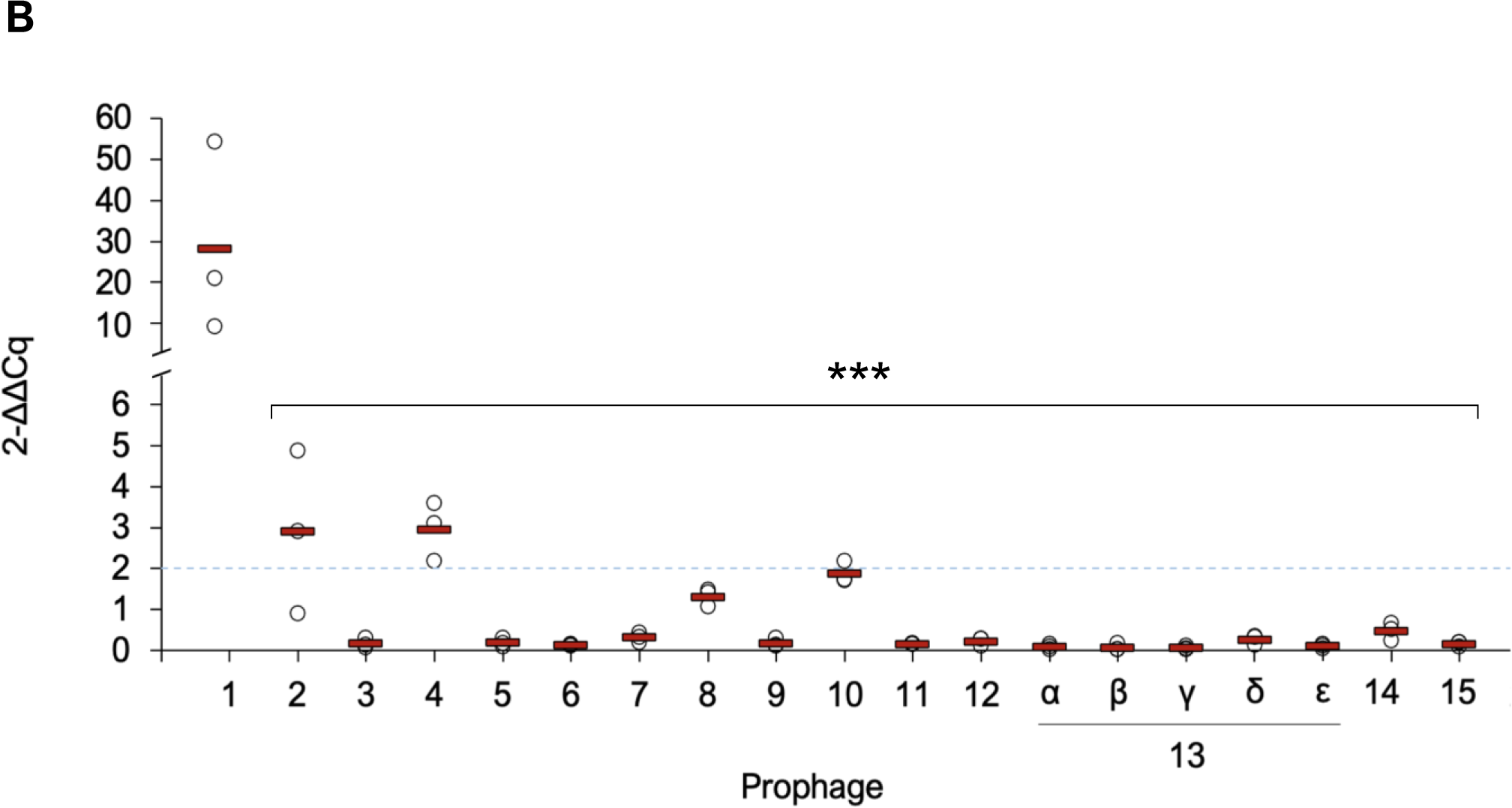
Basal release of *C. clostridioforme* LM41 prophages under non-inducing conditions. *C. clostridioforme* LM41 was diluted 1:50 from an overnight culture into standard FAB and grown for 24h under anaerobic conditions. Sterile filtered supernatants were subject to either no treatment or digestion with 10 μg/ml DNase I for 1.5h. 1 μl of DNase-treated or control supernatant was used as template for qPCR. *C. clostridioforme s10p* (equivalent to *rps*J) was used as the housekeeper and as a marker for the presence of bacterial DNA. **A.** Raw Cq values for the different target sequences. Data points are from 3 independent experiments with mean values shown as red bars. No template control Cq values for each target are shown by the dashed blue line. **B.** Fold-change in Cq of phage DNA in supernatant samples following DNase treatment to remove non-encapsidated DNA. The ΔCq values for all target samples were normalised using the ΔCq for the *s10p* gene (DNase treated Cq – untreated Cq), and fold changes were calculated using the 2^-ΔΔCq^ calculation. Data points are from 3 independent experiments with mean values shown as red bars. The dashed blue line indicates 2-fold change threshold for reference. Asterisks denote statistically significant differences between the mean 2^-ΔΔCq^ values for φ1 and the other phages, tested using a repeated measures ANOVA with Tukey post-hoc tests (no Greisenham correction), where *p* values ranged <0.0001-0.0002.

To visualise the phage particles spontaneously released into the culture supernatant, we DNase treated filtered culture supernatants of LM41, then NaCl-PEG 8000 precipitated the phage capsids, which were subsequently imaged using negative staining and transmission electron microscopy (TEM). Icosahedral particles with diameters in the range 64-67 nm were observed (Figure 5A&B), which is consistent with the dimensions of staphylococcal phages with similar genome sizes^33^. We also observed one example of a smaller-sized capsid, with a diameter of 39.4 nm (Figure 5C). Some virion particles appeared to have short protrusions (∼7-8 nm length) emanating from the capsid, raising the possibility that they are podoviruses (Figures 5A). The other structures observed were consistent with siphovirus or myovirus morphology, with the smaller diameter particle associated with what appears to be a large tail structure approximately 78.8 nm in length and 22.8 nm wide (Figure 5C). Though it is impossible to determine from the imaging analysis which LM41 phages are present in the sample, it is likely that the majority of the podovirus-like particles observed are LM41φ1 given that our previous experiment indicated that φ1 is the most abundant phage and is predicted to have podovirus morphology.

**Figure 5:**
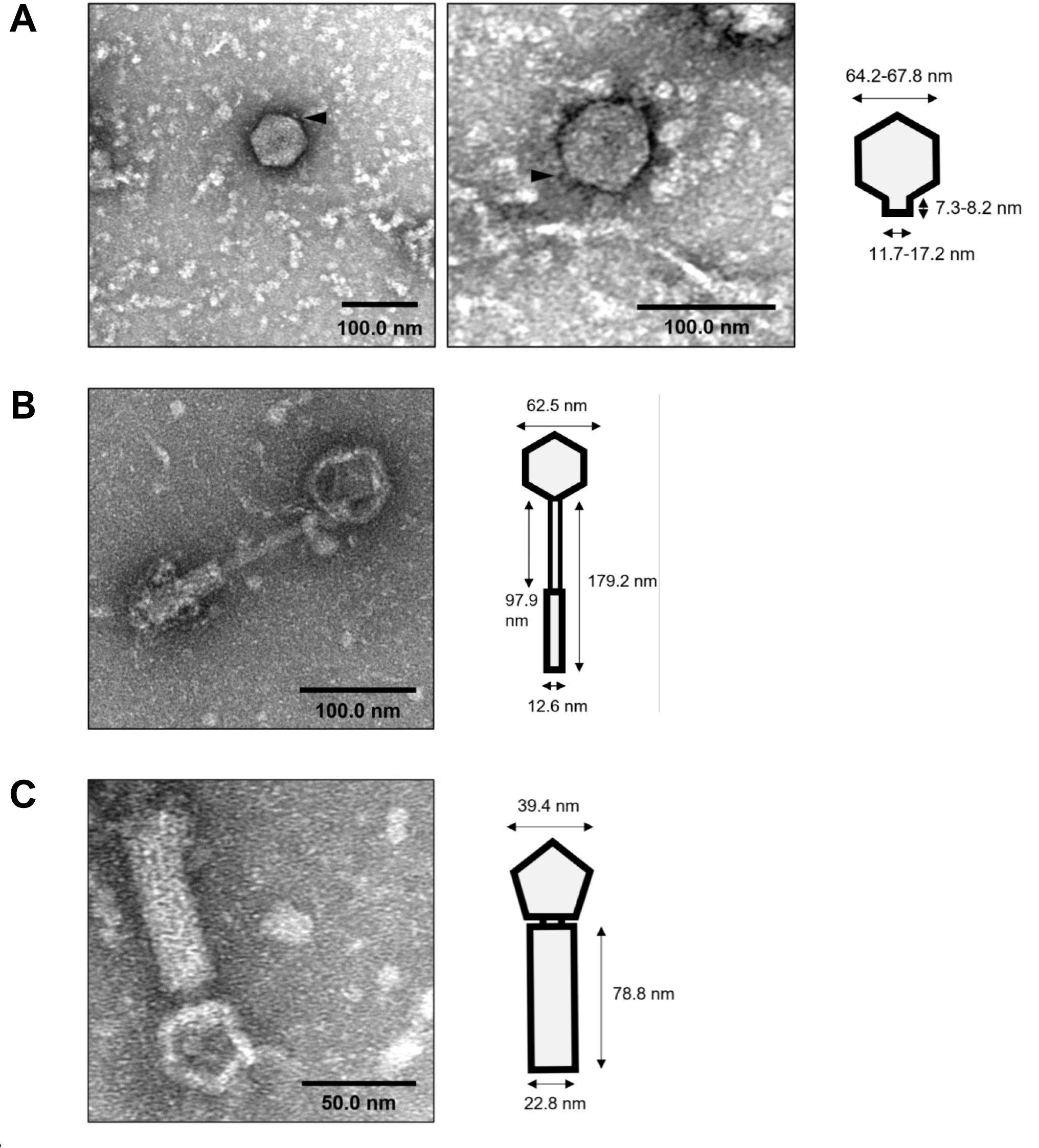
TEM examination of LM41 culture supernatant reveals phage particles. *C. clostridioforme* LM41 was diluted 1:50 from an overnight culture into standard FAB and grown for 24 h under anaerobic conditions. Sterile filtered supernatants were subject to digestion with 10 μg/ml DNase I for 1.5 h to remove non-encapsidated DNA, then concentrated following 10% PEG, 1 M NaCl precipitation. Samples were fixed and negatively stained with 0.5% (w/v) uranyl formate on copper-coated carbon grids, then imaged using a JEOL 1400 Flash TEM running at 80 kV. Particle dimensions for putative podoviruses (panel A) and tailed particles (panels B and C) are indicated on the schematics. Putative tail projections are indicated by black triangles (panel A).

### Accessory genes

We next sought to determine whether there was an obvious advantage to LM41 in maintaining so many prophage sequences. Bacteriophages often carry accessory genes that do not directly contribute to the lysogenic lifecycle but that may provide a benefit to the host bacterium by altering its phenotype in a process known as lysogenic conversion^34^. Importantly, accessory genes may also be retained as part of cryptic (defective) prophages^35^. We examined LM41φ1-15 for the presence of accessory genes that may confer some sort of benefit to the LM41 host cell. No products classically associated with bacterial morons (‘more-on’s), such as exotoxins or immune-evasion factors, were observed in any of the prophages encoded by LM41. This was not necessarily surprising, as *C. clostridioforme* is a member of the healthy gut microbiota and is not considered to be virulent. It should, however, be stated that many of the putative ORFs encoded by these prophages are predicted to be hypothetical proteins, so the presence of such factors cannot be definitively ruled out. We did note the presence of several potentially interesting proteins which can be broadly grouped as: restriction-modification (RM) system components; diversity-generating elements; hypothetical proteins with some similarity to large polyvalent proteins; phage defence factors; anti-phage defence factors; and proteins with possible roles in adaptation within the host intestine (Table 3).

**Table 3:**
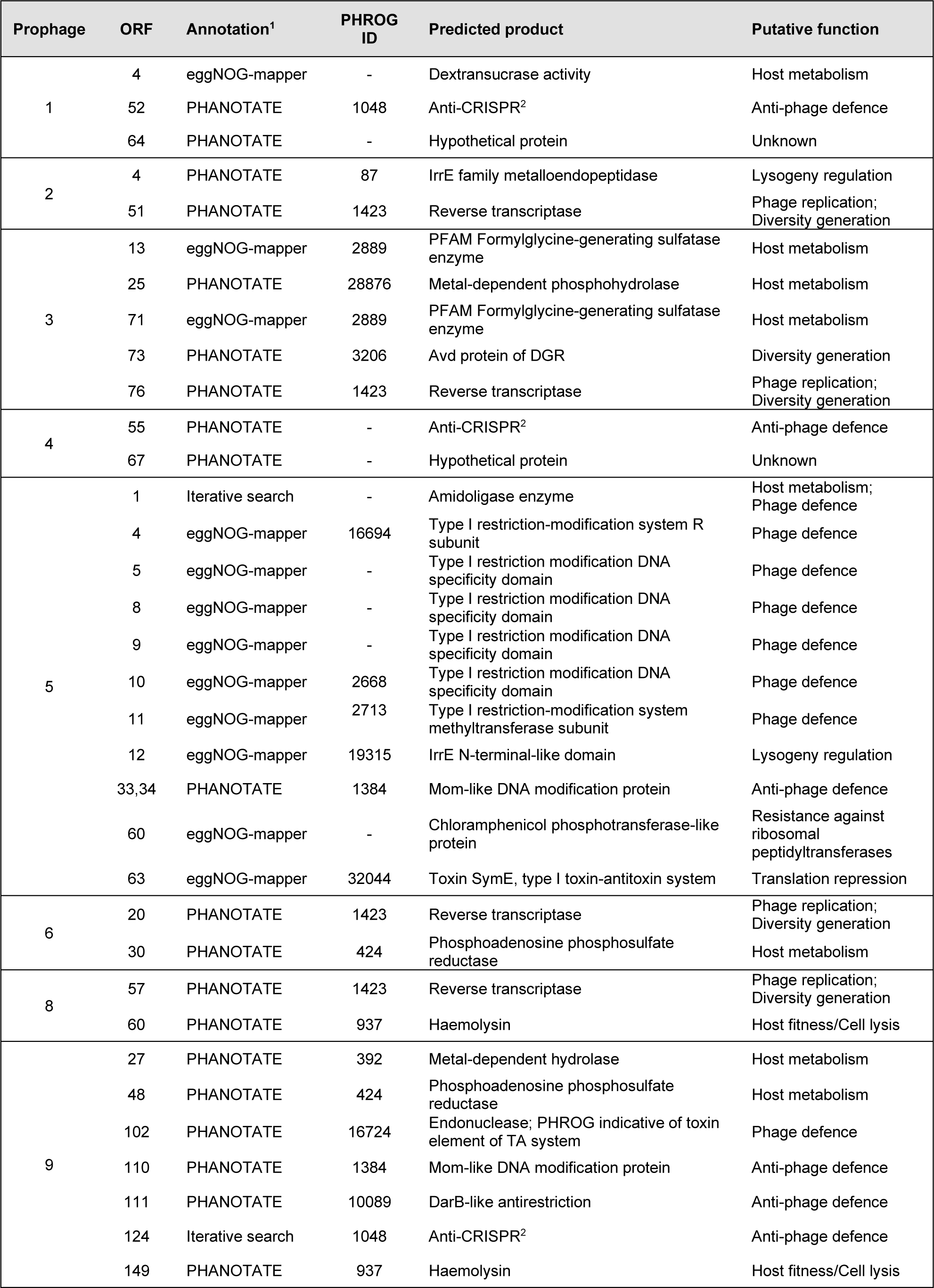
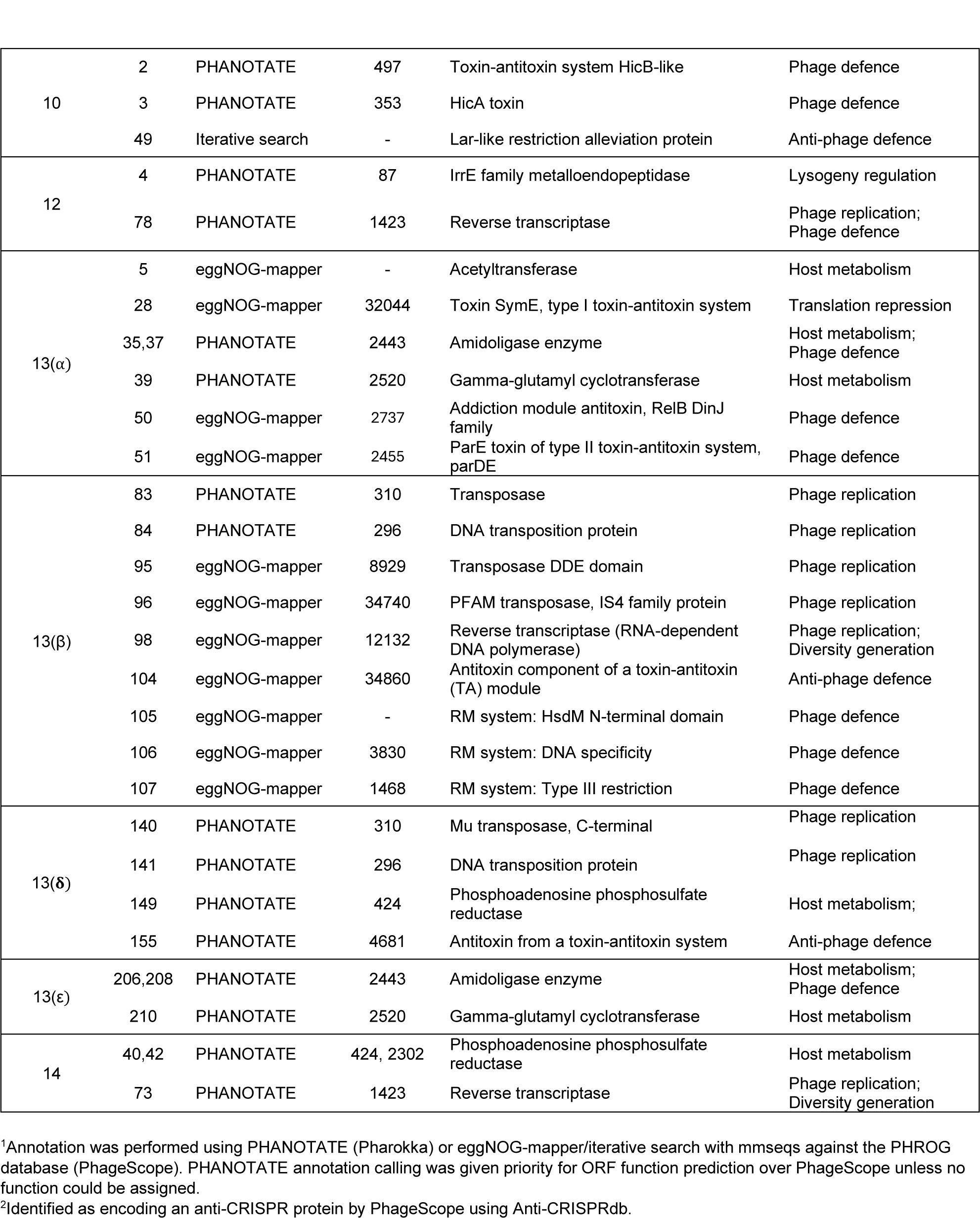
Prophage-encoded accessory Orfs of interest.

Proteins with roles in Type I RM were identified in LM41φ5, LM41φ13β and LM41φ13ε. Prophage LM41φ5 encodes a complete Type I RM system, encompassing three specificity subunits (*hsd*S), a DNA methyltransferase and a restriction subunit. LM41φ13β also encodes a complete Type I RM system comprising of HsdM (SAM-dependent DNA methyltransferase), HsdS (specificity subunit), and HsdR (endonuclease subunit R), where subunits HsdM and HsdR have high homology with similar proteins in *Ruminococcus sp.* (both 97.31% identity), while HsdS shares some similarity with a protein from *Anaerosporobacter faecicola* (60.22% identity). Interestingly, we also identified an ORF predicted to have limited similarity (61.11% identity) to the *E. coli* restriction alleviation protein, Lar (also known as RalR), in LM41φ10. Lar functions to modulate the activity of the *E. coli* K-12 host RM systems in order to protect the Rac prophage from destruction^36^.

In addition to the RM systems, we also noted the presence of a number of other factors associated with potential phage defence and anti-phage defence systems. Multiple different Toxin-Antitoxin (TA) systems were associated with the LM41 prophages: a HicB/HicA-type system identified in LM41φ10; a RelB/RelE-type system in LM41φ13α; and a SymE-like type I toxin in both LM41φ5 and LM41φ13α. Amidoligase enzymes were also identified in prophages φ5 and φ13α. In terms of anti-phage defence systems, anti-CRISPR systems were identified in φ1, φ2 and φ9, while a predicted TA system antitoxin was observed in φ13δ.

The presence of diversity-generating elements was also noted in multiple LM41 prophages. Group II intron reverse transcriptase/maturase proteins were identified in prophages LM41φ2 (Orf51), LM41φ6 (Orf20), and LM41φ12 (Orf78), while LM41φ3 is predicted to carry both a reverse transcriptase/maturase family protein (Orf76) and a homolog of *Bordetella* phage BPP-1 diversity-generating retroelement protein Avd (Orf73), which in BPP-1 facilitates sequence variation in target protein genes, enabling changes in host cell surface factors^37,38^.

Proteins with potential roles in influencing adaptation of the bacterial lysogen within the host intestinal environment were also observed. These include Orf60 of LM41φ8 and Orf149 of LM41φ9, which encode putative haemolysins, and factors influencing host metabolism, such as a gamma-glutamyl cyclotransferases in φ13α and φ13ε, an ORF with dextransucrase activity in φ1, and phosphoadenosine phosphosulfate reductases in φ13δ and φ14.

Finally, prophages LM41φ1 and LM41φ4 encode an unusually large (8 kb) ORF at their 3’ ends, which constitutes almost 20% of the phage genome sequence. Both ORFs are predicted to be hypothetical proteins, but have some limited similarity (53.84% ID) to large polyvalent protein-associated domain 3 from *Podoviridae* sp., suggesting that they could play a role in protecting and establishing the phage DNA when it enters a new host cell^39^.

## Discussion

*C. clostridioforme* LM41 has an atypically large genome for this species, with a strikingly high proportion of DNA attributed to mobile genetic elements^18^. This work sought to characterise the prophage sequences associated with this strain to determine whether they might contribute to its enhanced fitness in the dysbiotic gut. Interrogation of the LM41 genome sequence revealed poly-lysogeny: 15 prophage-derived sequences – comprising >9.6% of the bacterium’s 7.78 Mb genome – were observed, many with genomic organisation and size reminiscent of the well-characterised Gram positive staphylococcal siphovirus phages^40^. Four of the LM41 phages are predicted to encode all of the necessary modules for functionality, with a further nine phages requiring additional characterisation. We attempted to test functionality of the LM41 prophages experimentally using chemical induction, however the classical SOS-inducing antibiotics mitomycin C, norfloxacin and ciprofloxacin failed to induce bacterial lysis or significantly increase the quantity of encapsidated DNA released from LM41 compared to the untreated control, suggesting that LM41 prophages do not respond efficiently to this type of inducing signal. This observation is not necessarily surprising as work in *Clostridioides difficile* has shown that some prophages respond more effectively to fluoroquinolone antibiotic exposure than to the ‘gold-standard’ mitomycin C^41^. Furthermore, others have shown that in a variety of species of human gut bacteria, fewer than one quarter of prophages predicted to be functional using bioinformatics could be induced under experimental conditions^42^. This may suggest higher than expected rates of cryptic phage carriage in these bacteria or could mean that prophages in these species have different inducing signals to those from classically studied hosts such as *E. coli* and *S. aureus*. Arguably, it is likely that other prophage inducing signals occur in the gut environment given the lack of potent DNA damaging agents typically present in physiological habitats, and recent work has shown that in *Vibrio spp*., prophage-encoded transcription factors can activate small proteins which induce their prophage in an SOS-independent manner^43^, while *S. aureus* prophage phiMBL3 can be induced independently of the SOS response by a pyocyanin metabolite from *Pseudomonas aeruginosa*^44^. Accordingly, further work is necessary to screen a variety of inducing agents against LM41 prophages before they can be conclusively determined to be functional or defective.

Within lysogenic populations, spontaneous prophage induction can occur in a small proportion of cells, leading to release of low titres of phage into the surrounding environment^45^. Molecular examination of LM41 culture supernatants confirmed that LM41φ1, φ4 and φ10 particles are spontaneously released, supporting our prediction of these prophages as functional. φ2 was also detected, suggesting that this phage is functional despite scoring <100 for completeness. TEM imaging showed that spontaneously produced particles are predominantly podoviruses, though observation of other putative phage particles with longer tails indicates diversity in the morphological characteristics of LM41 phages. We also observed diversity among the lysogeny control systems utilised by the different prophages, suggesting the existence of a diverse community of phages within the *Lachnospiraceae* that can employ different mechanisms in order to maintain their latent state within their host bacterium.

Three regions were also observed that contained phage remnants to varying degrees, with the defects present predicted to abolish the ability of these phages to excise, replicate, and/or package efficiently. LM41φ13 was revealed to be not just one phage, but a 136 kb region of phage remnants, presumably derived from excessive or uncontrolled recombination events. A similar Mu-type phage is present in the *C. clostridioforme* LM41 relative *Lachnoclostridium* sp. YL32 (Accession: CP015399), where two copies of the 35.8 kb phage sequence are arranged divergently at genome locations 3,363,626-3,399,467 bp and 3,591,511-3,627,351 bp, with each of these sequences displaying high similarity (95.82% ID, 89% cover, E-value 0.0) to the δ region of LM41φ13, suggesting the potential for a common ancestor. It is unclear as to how and why the LM41φ13 region became so variable in LM41. In contrast to many well-characterised lysogenic phages, transposable phages do not excise out of the chromosome in order to proliferate^46^. In the case of the archetypal phage Mu, the integrated phage replicates by looping the bacterial chromosome and cleaving the DNA, enabling the formation of Shapiro intermediate structures whereby the prophage sequence is duplicated and integrated into new sites on the bacterial chromosome at random, in a mechanism similar to a transposon^46^. We can see no obvious explanation for the hypervariability observed in region LM41φ13, however, given the apparent propensity for DNA acquisition by strain LM41, it is possible that this strain has lost some of the mechanisms required for maintaining fidelity in DNA recombination and repair, resulting in the loss of intact phage regions. Additional work will be required to evaluate this theory further.

Given the quantity of prophage DNA carried by LM41, we hypothesised that one or more of the resident prophages contributes to the fitness of the host bacteria in the dysbiotic intestine. Examination of the prophage sequences for the presence of morons (accessory genes with functions not linked to lysogeny) revealed no obvious candidates for the enhanced fitness displayed by LM41 in the gut environment. We did, however, find that the LM41 prophages carry a number of potentially interesting genes, including those with roles in phage defence and anti-phage defence. Defence systems include a variety of DNA methyltransferases, a number of specificity subunits, and two complete Type I RM systems for the modification of DNA, presumably to aid phage defence against degradation by the host bacterium’s RM systems. Plasmid carriage of orphan HsdS (specificity) subunits that can interact with chromosomally-encoded HsdM (methylation) and HsdR (restriction) subunits has been described in *Lactococcus lactis*, creating a molecular expansion pack for the host cell Type I RM repertoire without requiring carriage of a full HsdMSR system^47^. It is tempting to speculate that a similar combinational system utilising phage-encoded specificity or methyltransferase subunits with native restriction and/or methylation components may contribute to the ability of LM41 to accept foreign DNA if it can be recognised and methylated by these enzymes prior to destruction by the host cell’s restriction systems, possibly lending some explanation as to why this strain appears to have gained so much horizontally-acquired DNA compared to its most closely related strains. Further to this, we observed a protein in LM41φ10 with limited similarity to the *E. coli* restriction alleviation protein, Lar, which has a role in modulating the activity of *E. coli*-encoded RM systems to protect prophage DNA^36^. It is not impossible that the Lar-like protein of LM41φ10 exerts global impacts upon its host organism, and that this could function synergistically with the other phage-encoded RM components to retain foreign DNA in LM41. In order to test this theory, a phage-cured strain of LM41 would need to be generated, and its ability to accept exogenous DNA compared with the parental LM41 and with variants carrying defined combinations of prophages. Unfortunately, given the paucity of genetic tools to permit manipulation of this organism, such experiments are not currently possible.

Other putative defence systems include TA systems, which can facilitate phage defence at the population level, inducing processes such as abortive infection following infection of the host cell or by inhibiting virion formation^48,49^. We also observed factors with roles in modifying the host cell surface to prevent superinfection of lysogens, potentially acting similarly to the amidoligase of *E. coli* phage phiEco32 which modifies cell wall receptors to prevent adsorption by competing phages^50^. As bacteria and their phages are engaged in a constant arms race, evolution of anti-phage defence systems on the part of the phage is necessary to overcome bacterial defences. LM41 prophages encode anti-CRISPR proteins and carry solitary TA system antitoxin components with potential roles in subverting phage defence systems. It is currently unclear whether these antitoxins are part of degenerate TA systems or whether these proteins could act as anti-phage defence systems by enabling the phages to counter toxins from other host- or phage-encoded TA systems.

Group II reverse transcriptase/maturase proteins were also detected in a number of prophages. The role of these proteins for phage or host cell biology is unclear. Indeed, it is possible that these elements have been acquired elsewhere and have become integrated within the prophage sequences, as seems likely in the case of LM41φ6 where the putative helicase ORF has been interrupted by the insertion of a putative group II reverse transcriptase/maturase protein. This hypothesis is further supported by the fact that 29 LtrA group II intron sequences are present throughout the LM41 genome, suggesting that these are a feature of the host rather than the phages.

Finally, we detected two putative haemolysin proteins carried by prophages φ8 and φ9. It is possible that these are misannotations, as φ9 Orf149 shows high homology to CHAP domain-containing protein from *Enterocloster sp.* (BLASTp: 99% query cover, E-value 0.0, 98.45% ID to accession WP_256170368.1) which is predicted to have a role in peptidoglycan hydrolysis, suggesting a role in phage-mediated cell lysis. However, if these proteins are indeed haemolysins, they could potentially provide LM41 with an advantage in the gut, perhaps by scavenging iron from the host via haemolysis. The ability of LM41 to lyse erythrocytes could be evaluated *in vitro* to examine whether LM41 has the potential to participate in nutrient acquisition in this way.

Phages are the most numerous entities within the gut microbiome^51^, and yet the phages associated with members of the microbiota remain poorly characterised. Here, we have identified an interesting example of poly-lysogeny in *C. clostridioforme* strain LM41 and have utilised bioinformatic tools and experimental approaches to offer insight into some of the characteristics of these phages, shedding light on their potential impact upon their host bacterium.

## Author contributions

S.H. and D.W. designed the study and obtained funding. S.H. wrote the manuscript. S.H. and A.M. performed the experiments. S.H., A.M. and D.W. performed the analysis. K.T. assisted with experiments and performed the statistical analysis. M.M. prepared the samples and performed TEM imaging.

## Acknowledgements

The authors thank Ester Serrano for helpful advice and assistance in preparing phage samples for TEM.

## Supplementary Figures

**Supplementary Figure 1:**
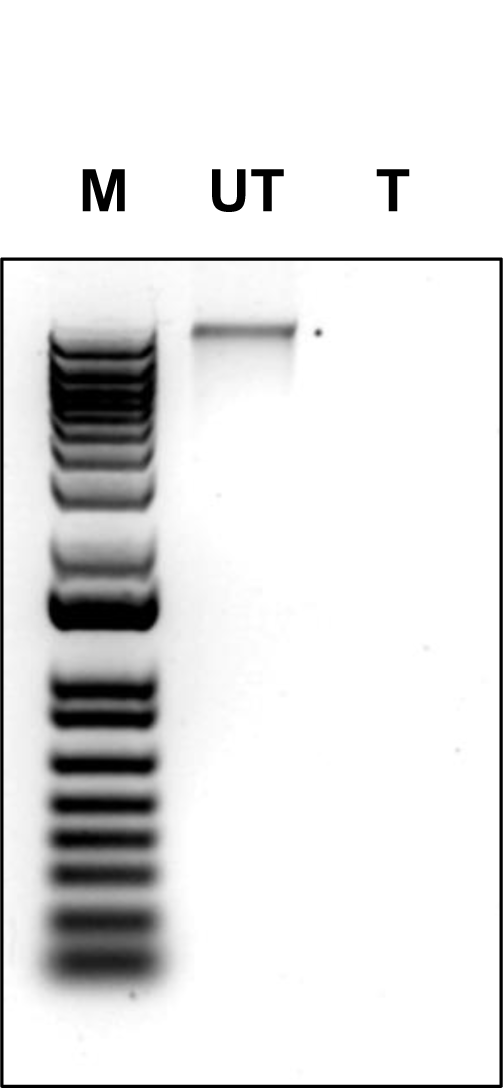
Confirmation of DNase activity against *C. clostridioforme* LM41 genomic DNA. 200 ng of *C. clostridioforme* LM41 genomic DNA was treated with (T) or without (UT) 10 μg/ml DNase I for 30 min at 37°C in parallel with culture supernatant sample digests (Figures 3 and 4) to confirm enzymatic activity. Samples were mixed with 10X loading dye and loaded on a 1% (w/v) TAE gel to run for 1.5 h at 120 V. 250 ng of 1 KB Plus DNA ladder (Invitrogen) was also loaded (M). The image shown is representative for each independent experiment performed.

